# Pseudogene-mediated gene conversion after CRISPR-Cas9 editing demonstrated by partial *CD33* conversion with *SIGLEC22P*

**DOI:** 10.1101/2021.04.20.440641

**Authors:** Benjamin C. Shaw, Steven Estus

## Abstract

Although gene editing workflows typically consider the possibility of off-target editing, pseudogene-directed homology repair has not, to our knowledge, been reported previously. Here, we employed a CRISPR-Cas9 strategy for targeted excision of exon 2 in *CD33* in U937 human monocyte cell line. Candidate clonal cell lines were screened by using a clinically relevant antibody known to label the IgV domain encoded by exon 2 (P67.6, gemtuzumab). In addition to the anticipated deletion of exon 2, we also found unexpected P67.6-negative cell lines which had apparently retained CD33 exon 2. Sequencing revealed that these lines underwent gene conversion from the nearby SIGLEC22P pseudogene during homology repair that resulted in three missense mutations relative to CD33. Ectopic expression studies confirmed that the P67.6 epitope is dependent upon these amino acids. In summation, we report that pseudogene-directed homology repair can lead to aberrant CRISPR gene editing.

## Introduction

The CRISPR-Cas9 system has revolutionized gene editing.^1^ In this process, a single-stranded guide RNA (sgRNA) directs Cas9 endonuclease to cleave DNA at a sequence-specific site. The DNA cleavage results in a double stranded DNA break (DSB) that is repaired by either homology-directed repair (HDR) or nonhomologous end-joining (NHEJ). The former results in targeted integration of DNA sequence while the latter typically results in gene disruption through the introduction of insertions or deletions (indels). To generate a targeted knock-in or knock-out, an HDR template of exogenous DNA is often supplied as part of the process.^2^ Alternatively, endogenous HDR templates have been described, including *HBD* sequence being incorporated into *HBB* and sequence from one allele of *HPRT* being incorporated into the other allele.^3, 4^ However, to our knowledge, HDR directed by a pseudogene has not been previously reported.

CD33 genetic variants, including rs124594919, have been associated with reduced risk of Alzheimer’s Disease (AD) in genome-wide studies.^5–7^ We and others subsequently identified rs12459419 as a functional SNP that increases the proportion of CD33 lacking exon 2 (D2-CD33).^8–12^ This exon encodes the ligand-binding IgV domain of this member of the sialic acid-binding immunoglobulin-type lectin (SIGLEC) family.^13^ Hence, while the extracellular portion of CD33 normally includes an IgV and IgC2 domain, D2-CD33 encodes a protein with only the IgC2 domain.^14^ CD33 inhibits microglial activity through its immunomodulatory tyrosine inhibitory motif (ITIM) and ITIM-like domains, which recruit protein tyrosine phosphatases, SHP1 and SHP2, to impact intracellular calcium flux, phagocytosis, and microglial migration.^9, 11, 12, 15–20^

Given that the AD-protective rs12459419 increases D2-CD33 at the expense of CD33, the prevailing theoretical mechanism has been that rs12459419 reduces AD risk through decreased CD33 function. However, recent findings that a *bona fide* loss of function indel, rs201074739, is not associated with AD risk, has led to this hypothesis being revised to suggest that rs12459419 and its related D2-CD33 isoform represent a gain of function.^13, 21, 22^ The gain-of-function mechanism and localization of D2-CD33 protein remain heavily debated. ^8, 9 11, 13, 17, 20–25^

Here, we sought to generate a model of physiologic D2-CD33 expression by using CRISPR-Cas9 to excise *CD33* exon 2 in the U937 human monocyte cell line. During these experiments, we identified a subset of cells which apparently underwent HDR directed by the *SIGLEC22P* pseudogene, located 13.5 kb away from *CD33*. Although the *SIGLEC22P* pseudogene shares approximately 87% identity over 1800 bp with *CD33*, this gene conversion was detected because three nucleotides in SIGLEC22P differ from those within the targeted *CD33* exon 2 and result in three missense amino acids in CD33, including p.N20K, p.F21I, and p.W22R. Hence, we report pseudogene directed gene conversion as a mechanism for unanticipated CRISPR mutations.

## Results

### CRISPR-Cas9-mediated CD33 Exon 2 deletion leads to loss of P67.6 epitope

To generate an *in vitro* model of D2-CD33, we targeted exon 2 for deletion by using guide RNAs corresponding to sequences in the flanking introns as previously described.^25^ Cells were transfected, maintained for two weeks, and sorted according to Figure 1. Live cells were gated by light scatter (Figure 1A), singlet events identified (Figure 1B), and sorted into separate tubes based on CD33 phenotypes (Figure 1C) with unstained cells shown for reference (Figure 1D). CD33 immunophenotype was determined with antibodies P67.6 and HIM3-4, which target epitopes in IgV and IgC2 that are encoded by exon 2 and exon 3, respectively. CD33 domains targeted by each antibody used in this study are shown in Table 1. We found that, of the 396,789 sorted cells, 91.7% fell outside of the unedited cell gate, and we presume these cells contain a CRISPR-mediated change in *CD33* sequence. Clonal cell lines were established from bulk collection of the gates drawn in Figure 1C. These cell lines were subsequently re-examined by flow cytometry with the same P67.6 and HIM3-4 antibodies. While unedited cells showed robust labeling by both HIM3-4 and P67.6 (Figure 2A), edited cell lines showed strong labeling by HIM3-4 but not P67.6 (Figure 2B), or no labeling by either HIM3-4 or P67.6 (Figure 2C). Data are representative of three independently established cell lines for each phenotype. Since D2-CD33 protein is not readily apparent on the cell surface^8, 21, 23–26^, we expected that the latter cells (Figure 2C) were candidates for exon 2 excision, which was confirmed by a PCR product of the appropriate size (Figure 2D, right) and by sequencing. However, cell lines with robust cell surface HIM3-4 but not P67.6 labeling were unexpected. Screening by the size of the PCR amplicon with primers corresponding to exon 1 and exon 3 suggested that exon 2 was still present (Figure 2D, middle). Sequencing of this PCR fragment revealed that the HIM3-4^+^ P67.6^-^ clones contained three apparent single nucleotide polymorphisms (SNPs) in exon 2 compared to the unedited, wild-type (WT-CD33) U937 cell line. These SNPs have an identical minor allele frequency (MAF) = 9.86 × 10^-5^ and are indexed as rs3987761, rs3987760, and rs35814802.^27^ Introduction of these SNPs results in changes in three consecutive amino acids (p.N20K, p.F21I, p.W22R) which we refer to as KIR-CD33. The nonsynonymous amino acids are the 4-6^th^ amino-terminal residues of the mature protein. EMBOSS and PSORT II predict cell surface localization for KIR-CD33 and no change in the signal peptide cleavage site.^28, 29^ Consistent with typical cell-surface localization, HIM3-4 labeled both CD33 and KIR-CD33 in a similar fashion (Figure 2A-B). Notably, total *CD33* gene expression and exon 2 splicing in KIR-CD33 cells does not differ from that of unedited cells (Figure 2E-F). CD33 expression was increased in D2-CD33 cells (Figure 2E) which exclusively express the *D2-CD33* isoform (Figure 2F).

**Figure 1.**
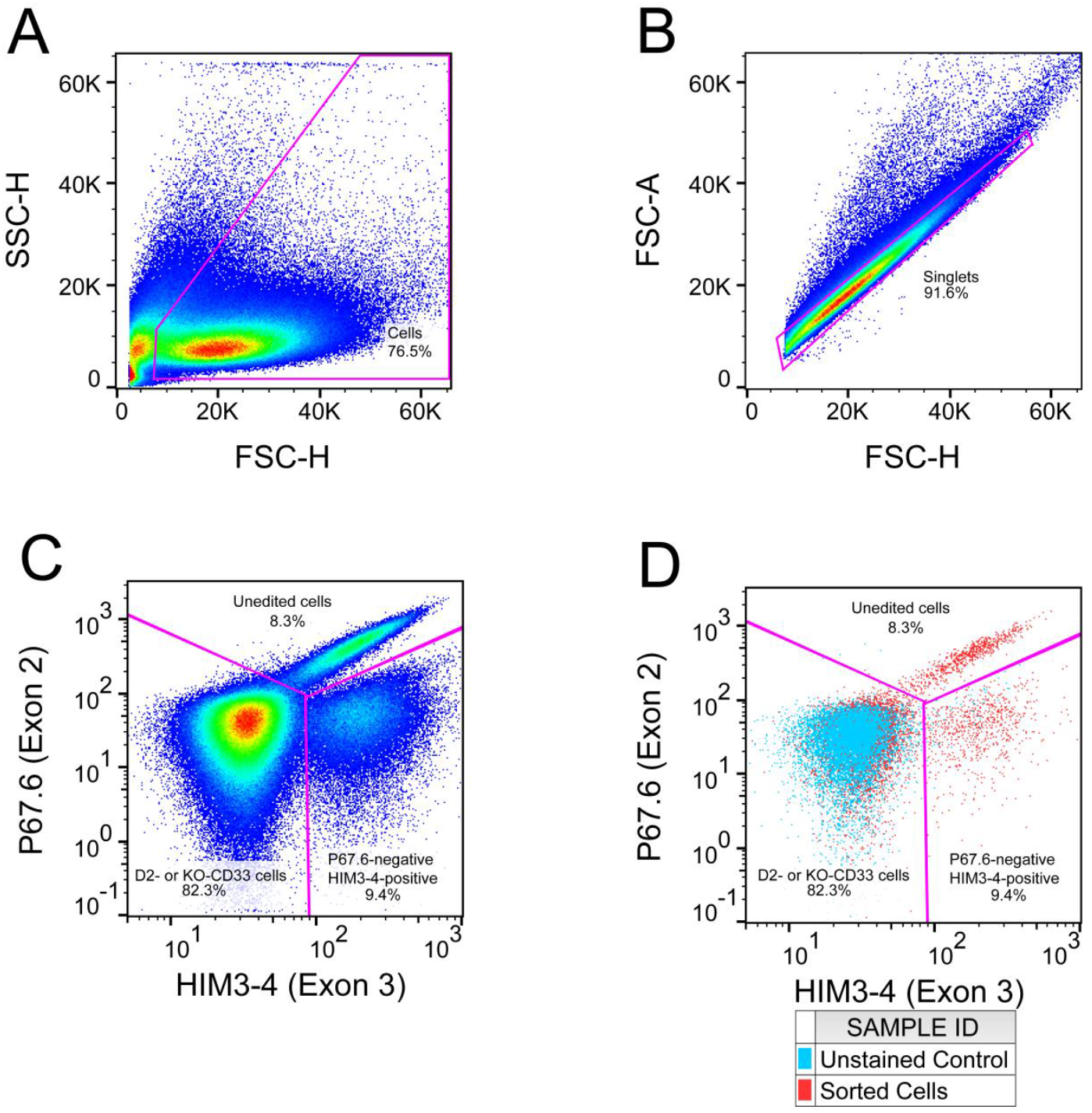
Sorting CRISPR-Cas9 *CD33* exon 2-edited cells reveals to two cell surface phenotypes. Gates are labeled in each panel and correspond to the following panel, with percent of parent gate shown. (A) Forward scatter (FSC) by side scatter gating was used to gate out dead cells (FSC^low^ SSC^high^). (B) Singlet events were selected along the FSC-Area by FSC-Height diagonal. (C) Populations were identified as either unedited (HIM3-4^+^ P67.6^+^), potential D2-CD33 or KO-CD33 (HIM3-4^-^ P67.6^-^), or unexpected (HIM3-4^+^ P67.6^-^). (D) Unstained control (blue) shown on top of the sorted cells (red) for reference. This unexpected population occurred at approximately a 1-to-9 frequency with respect to the D2- or KO-CD33 cells, or 10% of all edited cells.

**Figure 2.**
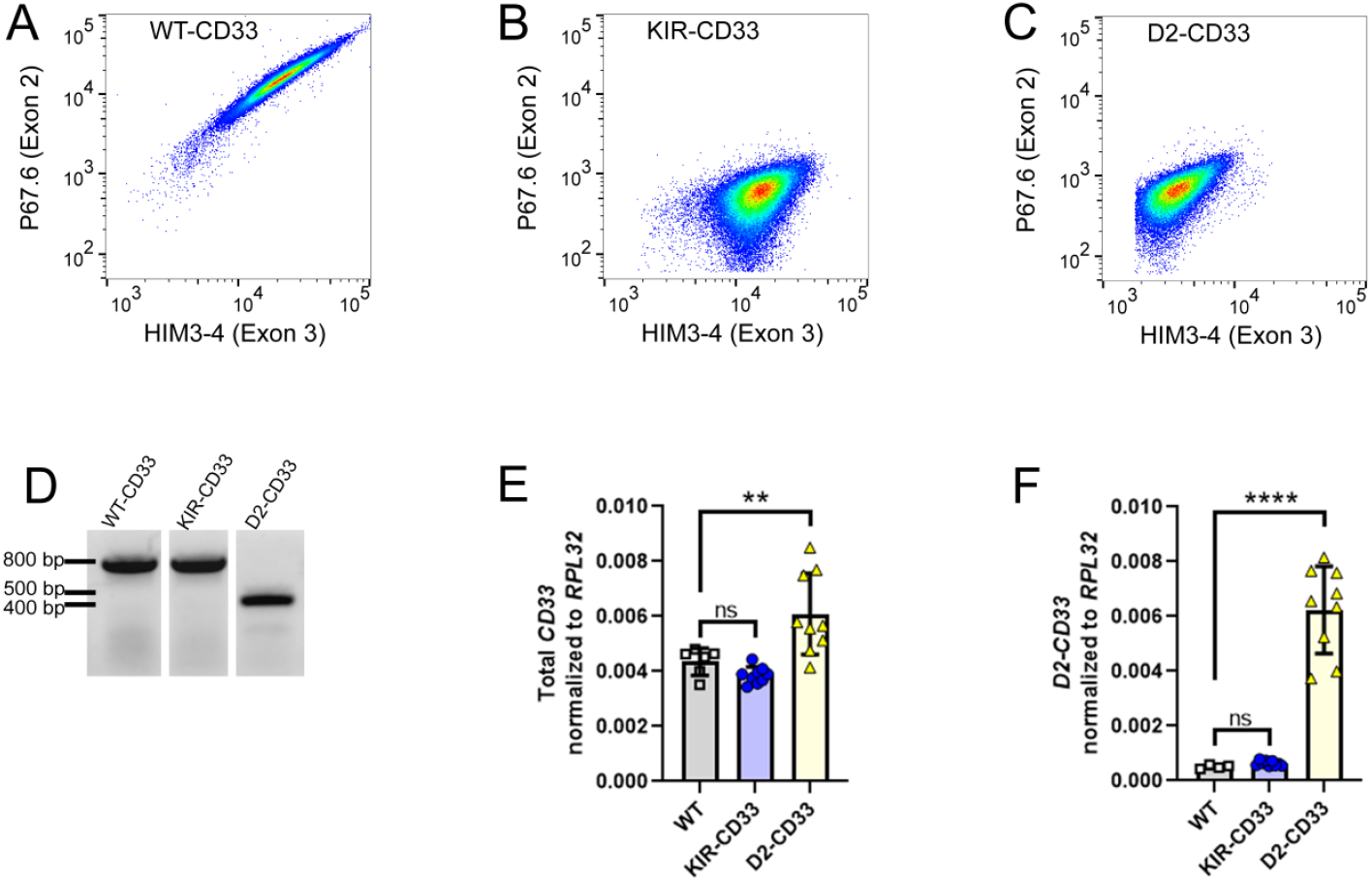
CRISPR-Cas9 editing of *CD33* exon 2 leads to loss of P67.6 epitope. (A) Unedited U937 cells display robust P67.6 and HIM3-4 labeling. (B) Edited U937 clone that is robustly labeled by HIM3-4 but not P67.6. (C) Edited U937 clone that is not labeled by either HIM3-4 or P67.6. The depicted results in B-C are representative of at least three clonal cell populations established for each phenotype. (D) Genomic DNA PCR of *CD33* exon 1 to exon 3 of the above cell lines. PCR products at 789 bp and 428 bp correspond to the expected sizes for the presence and absence of exon 2, respectively. (E) *KIR-CD33* mutations do not affect total *CD33* gene expression as determined by qPCR, but removal of exon 2 increases total *CD33* gene expression by 39.8%. (F) *KIR-CD33* mutations do not affect splicing efficiency of exon 2 as determined by qPCR, but removal of exon 2 at the genomic level increases exon 1-exon 3 junction to 100% of total *CD33* expression. Data from E-F analyzed by one-way ANOVA followed by Dunnett’s multiple comparisons test to unedited control. ** p < 0.01; **** p < 0.0001.

**Table 1.** Antibodies used in this study.

| Clone | Conjugate | Vendor | Catalog # | Lot | Dilution/Use | CD33 domain targeted |
| --- | --- | --- | --- | --- | --- | --- |
| P67.6 | BV711 | BioLegend | 366624 | B302694 | 2.5 µg/mL FC | IgV |
| P67.6 | Unconjugated | BioLegend | 825601 | B258818 | 5 µg/mL IF | IgV |
| WM53 | BV711 | BioLegend | 303424 | B253729 | 5 µg/mL FC | IgV |
| WM53 | Unconjugated | BioLegend | 96281 | B274701 | 5 µg/mL IF | IgV |
| HIM3-4 | FITC | BioLegend | 303304 | B284834 | 20 µg/mL FC | IgC<sub>2</sub> |
| PWS44 | Unconjugated | Leica | NCL-L-CD33 | 6024275 | 2 µg/mL IF | IgC<sub>2</sub> |
| H110 | Unconjugated | Santa Cruz | sc-28811 | K2211 | 1 µg/mL IF | Intracellular |
| Goat anti-Mouse | AlexaFluorPlus 488 | Invitrogen | A32723 | TF266577 | 10 µg/mL IF |  |
| Goat anti-Rabbit F(ab)<sub>2</sub> | AlexaFluor 568 | Invitrogen | A21069 | 2087701 | 10 µg/mL IF |  |

IF: immunofluorescence, FC: flow cytometry

### Validation of the CRISPR-induced in-frame disruption of the P67.6 epitope

To rigorously demonstrate that the lack of P67.6 labeling was solely due to the KIR mutations, we introduced the KIR mutations into a previously described CD33 expression vector.^8^ CD33 and KIR-CD33 vectors were transfected into HEK293 cells, which do not naturally express CD33, and the cells processed for flow cytometry (Figure 3) and immunofluorescent confocal microscopy (Figure 4). For flow cytometry, cells were labeled with P67.6 or WM53, each of which was conjugated to the same fluorochrome to facilitate direct comparisons (Figure 3). Importantly, these antibodies both target exon 2, are both mouse IgG1κ and have comparable degrees of labeling. In the CD33 HEK293 cells, labeling with both WM53 and P67.6 correlated well with HIM3-4 (Figure 3A-B). However, in the KIR-CD33 HEK293 cells, labeling with WM53 but not P67.6 correlated with HIM3-4 labeling as P67.6 labeling was not apparent (Figure 3C-D). Further gating on the HIM3-4^+^ cells shows that WM53 labeling is not affected by the KIR-CD33 mutation (Figure 3E), while P67.6 labeling in KIR-CD33 cells is comparable to non-transfected control cells (Figure 3F). These results were confirmed with immunofluorescent confocal microscopy using an array of anti-CD33 antibodies (Figure 4). The *CD33*-transfected HEK293 cells showed consistent double-labeling between an antibody against a cytoplasmic epitope (H110) and either WM53, P67.6, or PWS44 (Figure 4A-C). For the KIR-CD33 cells, robust co-labeling was observed between H110 and WM53 or PWS44 (Figure 4D, F). However, P67.6 labeling of KIR-CD33 cells was not detected (Figure 4E). We thus conclude that the residues identified—p.N20, p.F21, and p.W22—are necessary for P67.6 binding and that these changes are the reason that this antibody failed to label the CRISPR-edited cells. Although P67.6 has been humanized and used clinically, prior studies have not mapped its epitope at this resolution.^17, 30, 31^

**Figure 3.**
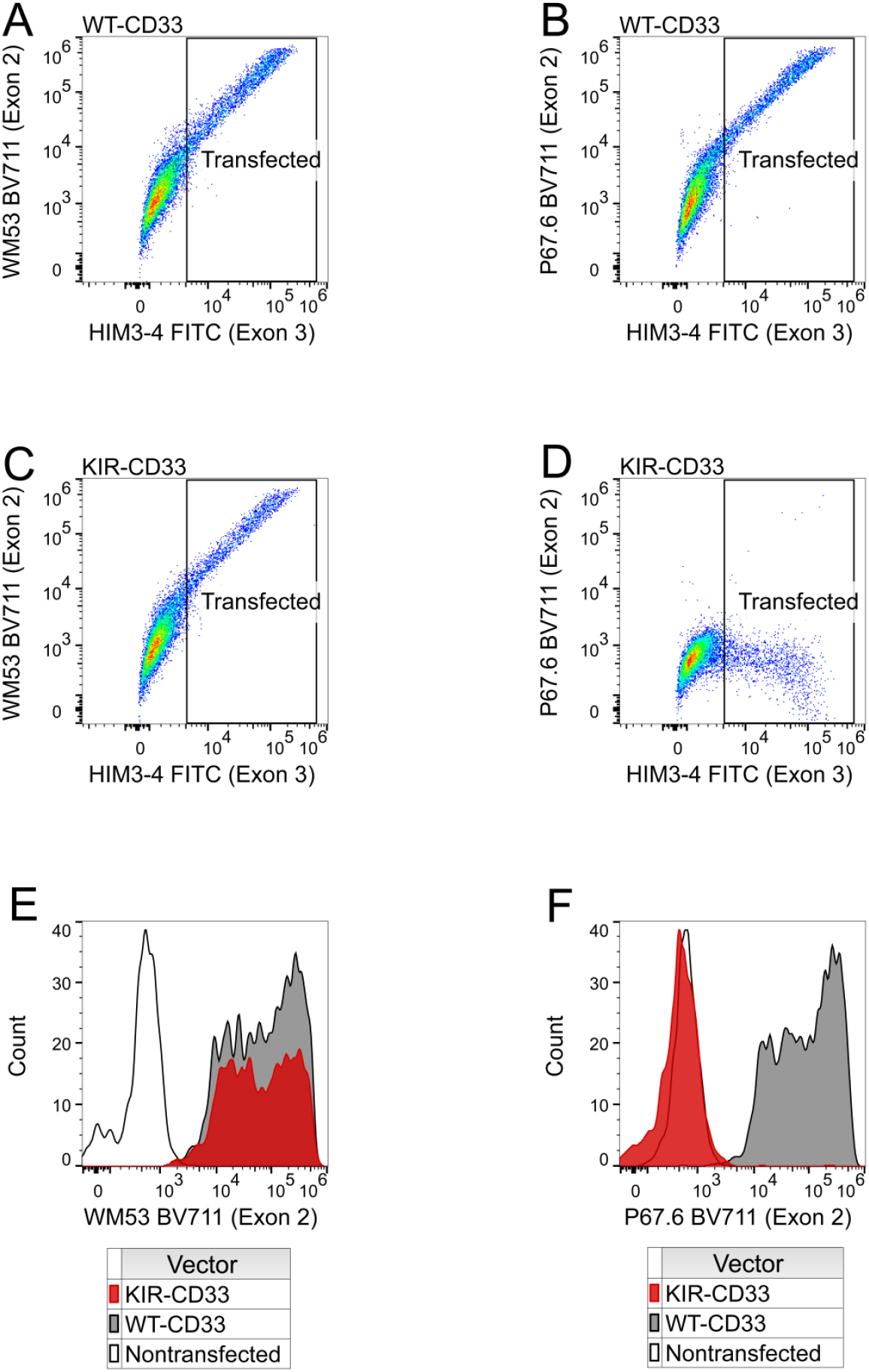
Loss of P67.6 epitope is in-frame and preserves cell surface expression. WT-CD33 (A, B) or KIR-CD33 (C, D) expressing HEK293 cells were labeled with HIM3-4 and either WM53 (A, C) or P67.6 (B, D). HIM3-4^+^ cells identified in the “Transfected” gate (A-D) were gated to show WM53 (E) or P67.6 (F) binding in transfected cells.

**Figure 4.**
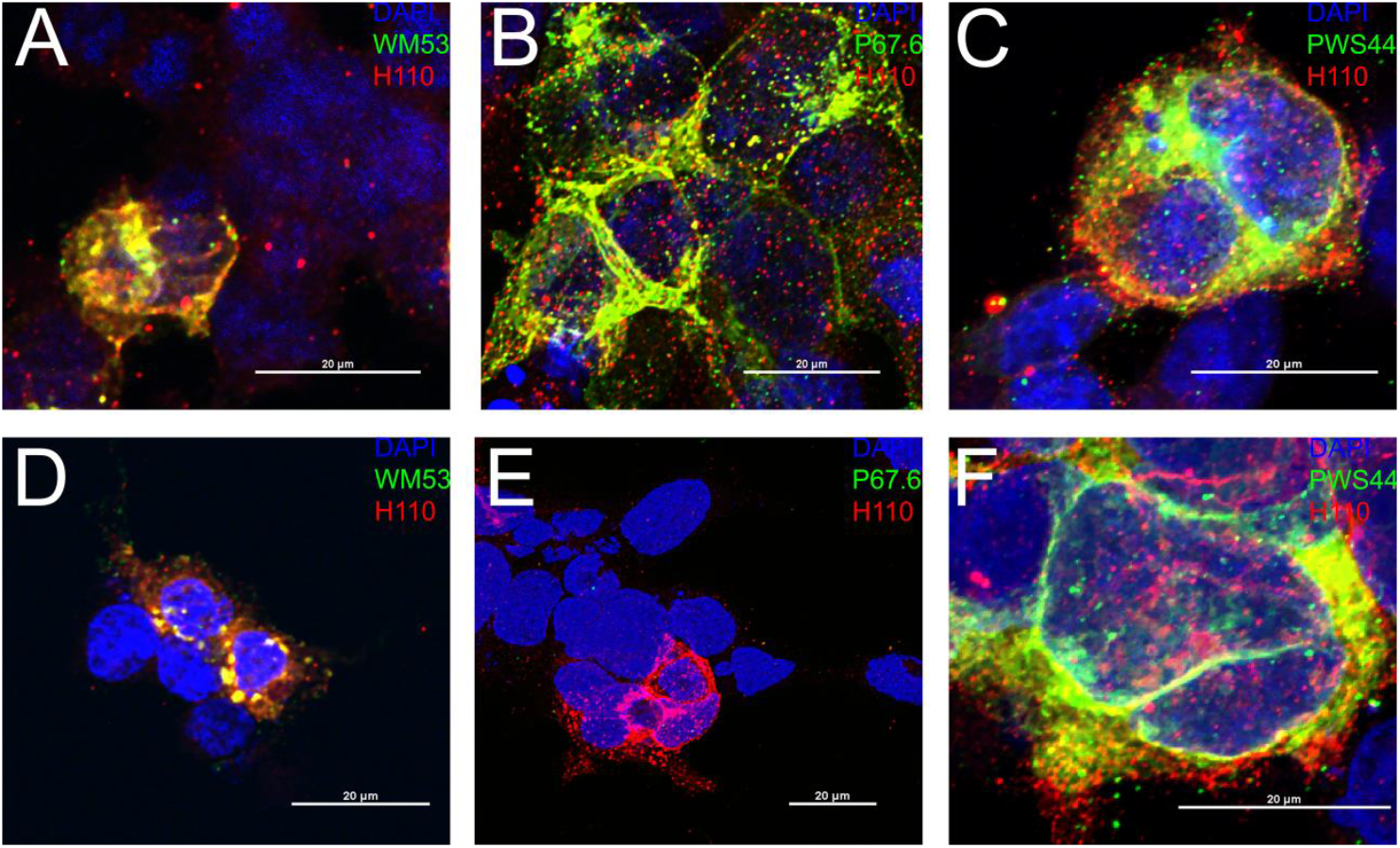
Loss of P67.6 epitope does not alter other common CD33 epitopes. WT-CD33 (A-C) or KIR-CD33 (D-F) HEK293 cells were labeled with H110 (red) and either WM53 (A, D; green), P67.6 (B, E; green), or PWS44 (C, F; green) and DAPI (blue).

### Identification of SIGLEC22P as a homology-directed repair template for CD33

The three nucleotide changes (Figure 5, red) are 23-27 bp away from the putative Cas9 cleavage site (Figure 5, scissors), were not consecutive, and were present and identical in each of the clones with the KIR-CD33 phenotype. Since this was unlikely due to chance, we hypothesized that this was due to HDR from elsewhere in the genome. A search of a 50 bp sequence centered on the KIR mutations revealed that this sequence occurs in the *SIGLEC22P* pseudogene which is located 13.5 kb away from *CD33*. Further investigation found an extended region of homology between *CD33* and *SIGLEC22P* that flanked the Cas9 cleavage site; indeed, the first 500 bp of the genes share 97% identity, including a 143 bp region of otherwise complete identity centered on the KIR mutations (Figure 5, underline). The *SIGLEC22P* pseudogene exon 2 contains 11 additional mutations as well as two intronic mutations relative to CD33. None of these mutations were detected in the CRISPR-edited cell lines, indicating the region used for repair was limited to, at most, the 143 bp region surrounding the KIR mutations near the Cas9-induced double-stranded break. We interpret these results as indicating that, upon double-strand breakage at the beginning of exon 2 in *CD33* (Figure 5, blue), the *SIGLEC22P* pseudogene was used as a repair template because of the strong sequence homology to *CD33* (Figure 5, underlined). In this process, three *SIGLEC22P*-specific mutations were introduced into *CD33*, resulting in missense mutations in three adjacent codons and thus KIR-CD33. This indicates an in-frame, ectopic gene conversion using pseudogene sequence. Two lines of evidence indicate that the KIR-CD33 cell lines were homozygous for the KIR mutation. First, the DNA sequence chromatogram showed only clear single peaks through the KIR sequence (Figure 5, chromatogram). Second, cell populations with an intermediate level of P67.6+ labeling were not detected (Figure 1C). Further, sequencing of the *SIGLEC22P* region in the KIR-CD33 clones revealed that the *CD33* sequence was not present, which we interpret to mean that *SIGLEC22P* was used as a repair template, rather than a crossover event.

**Figure 5.**
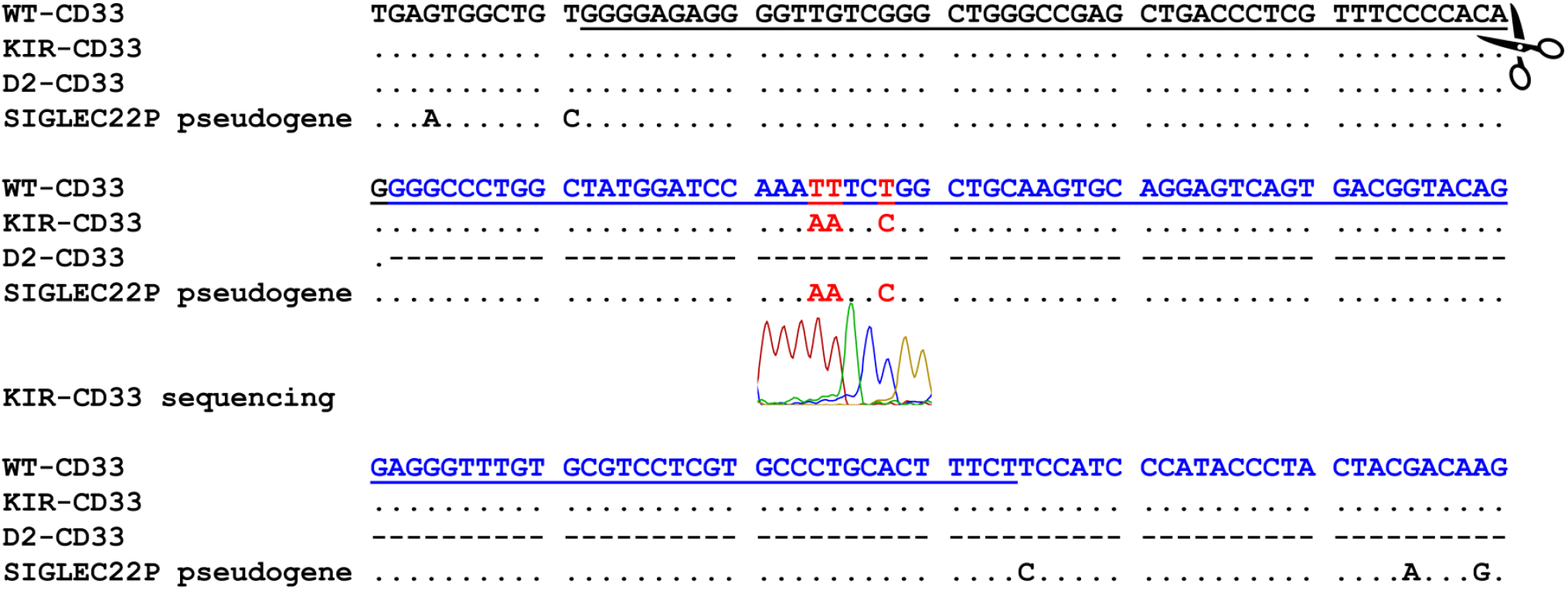
Alignment of cell line sequencing data. Unedited U937 cells at top, followed by KIR-CD33 sequence, D2-CD33 sequence, and reference *SIGLEC22P* sequence. Cas9 cleavage site marked by scissor icons. Mismatches from the unedited *CD33* sequence are denoted by either the differing base (KIR-CD33) or dash in the case of a gap (D2-CD33). Intron 1 in black, exon 2 in blue. The *SIGLEC22P* region used as a repair template is underlined. Mutations introduced into *CD33* in red. A representative post-PCR sequencing chromatogram from one clonal KIR-CD33 is shown at the mutation site with clear, single peaks demonstrating homozygosity.

## Discussion

We show here that the DSB repair pathways initiated after S. pyogenes Cas9 cleavage can lead to ectopic gene conversion from a pseudogene in a mitotic human cell line. This gene conversion resulted in an in-frame chimeric protein, wherein less than 150 bp of sequence from a nearby pseudogene replaced the targeted gene sequence (Figure 5). Indeed, this distance is consistent with meiotic gene conversion events observed by Jeffreys et al., who showed that gene conversion occurs through relatively short tracts with a mean length between 55-290 bp.^32^ Given that the *SIGLEC22P* locus in these KIR-CD33 cells lacks any detectable *CD33* sequence, we interpret this as further evidence of a gene conversion, rather than a mitotic crossover event. We speculate that this gene-conversion occurs in *trans*, i.e. from the intact chromosome, as a *cis*-mediated repair would more likely result in a gene fusion with intergenic deletion as has been previously reported.^33, 34^ This type of deletion event also occurs naturally, in the absence of Cas9 DSBs, as in the case of *SIGLEC14* deletions.^35^ Gene conversion in *trans* after CRISPR-induced DSBs has been demonstrated previously.^4^ We were surprised by the unexpectedly high frequency of conversion observed here; this pseudogene conversion occurred at approximately 1 in 10 edited cells. Exogenous double-stranded regions of homology as short as 58 bp have been used *in vitro* to introduce mutations through CRISPR-Cas9/HDR mechanisms.^2^ Coincidentally, this ectopic gene conversion disrupted the epitope of a well-validated antibody, P67.6 (Figure 6), known clinically as gemtuzumab. Using transiently transfected HEK293 cells expressing the chimeric protein, we demonstrated that these mutations are sufficient to abrogate P67.6 binding, providing the most precise epitope mapping to date of this clinically relevant antibody.

**Figure 6.**
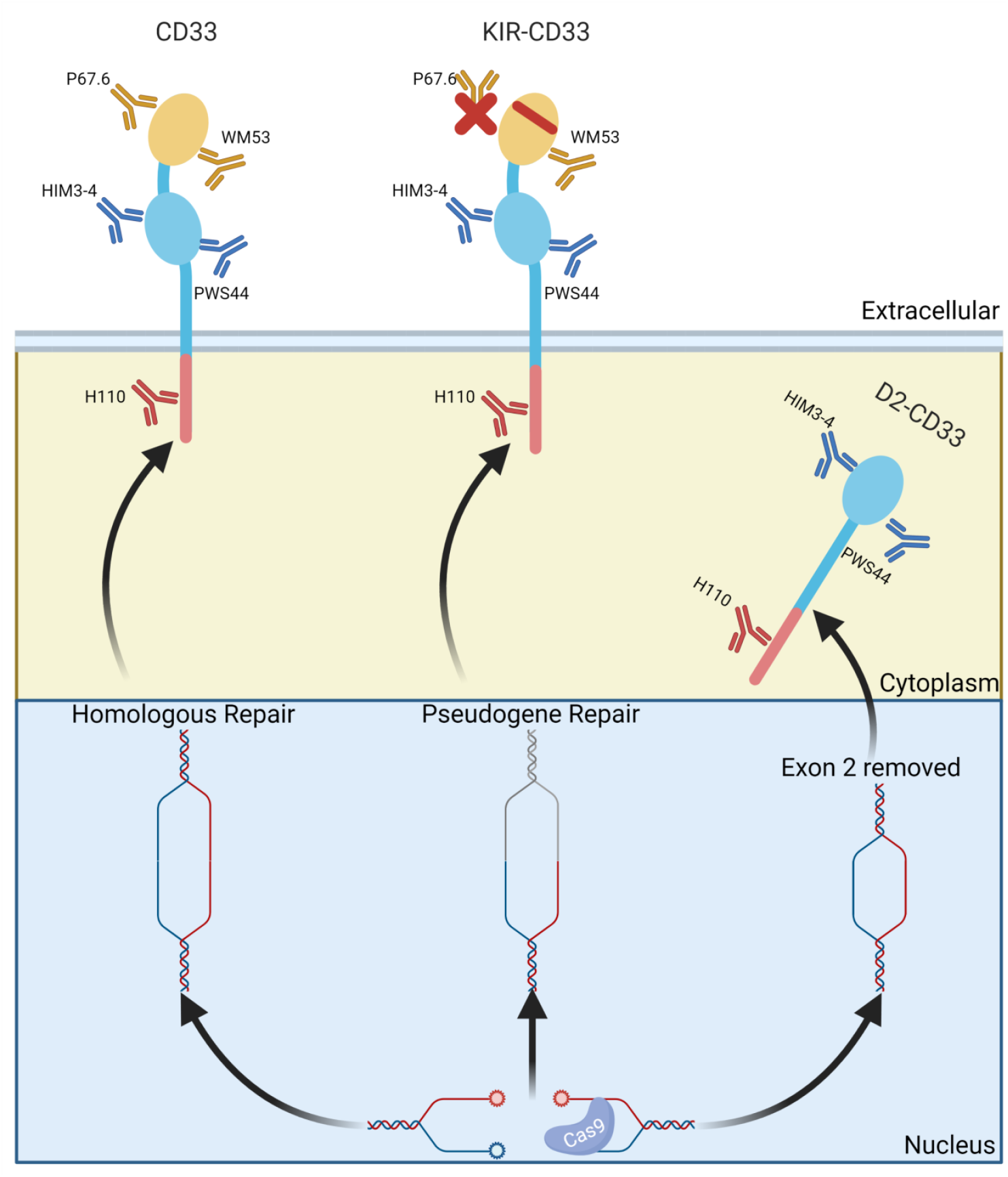
Model of pseudogene repair mechanism and anti-CD33 antibody binding sites. CD33 is normally a transmembrane, cell surface receptor with one IgV domain and one IgC2 domain. The KIR-CD33 mutation introduced by pseudogene directed-repair abrogates P67.6, but not WM53, binding. IgC2 domain antibodies HIM3-4 and PWS44, and the intracellular domain H110 antibody bind both CD33 and KIR-CD33. D2-CD33 is not readily apparent on the cell surface in CRISPR-Cas9 edited U937 cells, implying that under physiologic expression, the D2-CD33 protein is retained in an intracellular vesicle. Overall, pseudogene repair of a CRISPR-Cas9-targeted gene can disrupt the binding of well-validated antibodies without introducing a frameshift, and protein-level expression alone is not sufficient for knockout confirmation. Created with Biorender.

The KIR-CD33 mutations—rs3987761, rs3987760, rs35814802—are indexed in dbSNP and gnomAD with minor allele frequencies < 10^-4^, and roughly equivalent across populations.^27^ We considered the possibility that these mutations, while rare, occur naturally and are clinically relevant. The SIGLEC family of genes are undergoing rapid evolution in many species, including humans.^36^ This rapid evolution has resulted in the pseudogenization of many SIGLECs, including the *CD33* pseudogene *SIGLEC22P*. These same bases are indexed as SNPs in *SIGLEC22P* (rs997169007, rs1049597792, and rs1005338799), also have identical frequencies (MAF = 2.1 × 10^-4^) in the gnomAD database, and the major and minor alleles are the inverse of the KIR-CD33 SNPs.^27^ The region of homology identified here between *CD33* and *SIGLEC22P* is 143 bp, substantially shorter than the 50 bp reads upon which the gnomAD and dbSNP databases are built.^37^ While it is possible these are *bona fide* variants in *CD33*, the most likely explanation is that the *SIGLEC22* pseudogene sequences have been mapped incorrectly to *CD33* by algorithms. These missense SNPs are also not recorded in the BeatAML variant database, which records known AML-associated functional variants such as missense SNPs, further underscoring the low probability that these are true *CD33* variants.^38^

Off-target editing is a frequent concern in gene editing workflows, including CRISPR-Cas9. However, gene conversion is often overlooked as a potential confound. For instance, within *CD33*, off-target editing of *SIGLEC22P* resulting in a 14 kb deletion and subsequent *SIGLEC22P/CD33* gene fusion has been previously reported, which may have been the result of a mitotic crossover.^33, 34^ Neither report noted a gene conversion between *SIGLEC22P* and *CD33*. Mitotic crossover initiated by CRISPR-Cas9 cleavage has also been reported in the *HPRT* locus, resulting in a 36 kb crossover.^4^ Gene conversion via CRISPR-Cas9 between a 101 bp homologous region in *HBD* and *HBB* has also been reported; notably, these genes were also the first reported gene conversion event in humans.^3, 39^ To our knowledge, this is the first report of pseudogene-mediated gene conversion during the CRISPR-Cas9 editing process. Our results demonstrate the need for rigorous screening in studies which rely on gene editing, and analysis of the region flanking the expected cut site for homology and possible gene conversion. Since the human genome contains 8000-12000 pseudogenes and approximately 3400 genes in the human genome have known pseudogenes, pseudogene directed homology repair is a potentially considerable confound.^40, 41^ Approximately 84% of putative pseudogenes are estimated to be located on a different chromosome than their parent gene, while the remaining 16% have a mean intergenic distance of 1.8 Mb (median of 1.3 Mb).^42^ Whether the gene conversion described here occurs in cis or trans, or if this event is impacted by intergenic distance, is unclear. As pseudogenes are by definition nonfunctional, they may be overlooked as irrelevant during the design process. Given the pseudogene conversion described here, there is a clear need to analyze the sequence of the edit, rather than relying on gene or protein expression. This is especially relevant given that CRISPR-mediated *CD33* knockout allografts and autografts have been proposed as a potential AML treatment strategy in conjunction with chimeric antigen receptor (CAR) T cells.^33, 34^ In this strategy, autologous transplantation of *CD33*^null^ hematopoietic stem and progenitor cells (HSPCs) would reconstitute the myeloid system, while engraftment of CAR-T cells would provide long-term surveillance against any surviving CD33^+^ AML cells. Mitotic gene conversion, and presumably the herein described pseudogene-mediated gene conversion, has been demonstrated after CRISPR-Cas9 editing in primary human cells between *HBD* and *HBB* ^3^ Screening these HSPCs with the standard gemtuzumab antibody, however, could lead to engraftment of some KIR-CD33 cells as well.

While off-target editing of a nonfunctional pseudogene may have limited impact on downstream results, incorporating elements from a nonfunctional pseudogene into a target gene may lead to deleterious mutations that abrogate function of the target gene entirely, for instance the introduction of a premature stop codon. This is especially important to consider for designs which incorporate unedited cells which have undergone sorting and single cell cloning as a control. We also speculate that this pseudogene-mediated repair will reduce the efficiency of creating a gene disruption if the disruption site has homology in a pseudogene. The breakage site may be repaired by a pseudogene with complete identity at the DSB, requiring more clones to be screened to find a gene disruption. By coincidence, our initial screen for editing included an antibody which overlapped the KIR sequence. This gene conversion, at the protein level, is masked when using an alternative antibody (WM53) targeting the same domain and is not apparent by PCR alone. Combining the data from Figures 2D-F and 3C, one could reason that these cells were unedited as there are no apparent differences in PCR fragment size, gene expression, splicing, or cell surface protein expression, and thus incorrectly assume that sequencing is unnecessary. We conclude that, in addition to off-target Cas9-editing confounds, researchers should be aware of the potential for pseudogene-directed homology repair.

## Methods

### Cell Lines and Antibodies

U937 and HEK293 cell lines were obtained from American Type Culture Collection (ATCC). U937 cells were cultured in RPMI 1640 with HEPES (Gibco 22400-089) supplemented with 10% fetal bovine serum, defined (HyClone, GE Healthcare SH30070.03); 50 U/mL penicillin, 50 μg/mL streptomycin (Gibco 15070-063); and 2 μM L-glutamine (Gibco A2916801). HEK293 cells were cultured in EMEM, ATCC formulation (ATCC 30-2003) supplemented with 10% fetal bovine serum, defined (HyClone, GE Healthcare SH30070.03); 50 U/mL penicillin, 50 μg/mL streptomycin (Gibco 22400-089). Cells were maintained at 37°C in a 5% CO2 in air atmosphere. The U937 cell line has been reported as either diploid or triploid at chromosome 19 which contains *CD33*.^43, 44^Antibodies, concentrations, and CD33 domains targeted are shown in Table 1.

### CRISPR-Cas9 Gene Editing

CRISPR reagents were purchased from Integrated DNA Technologies (IDT). Single-guide RNAs (sgRNAs) and S. pyogenes Cas9 protein (IDT 1081059) were incubated at a 1:1 molar ratio (0.5 nmol each) at room temperature for 10 minutes to form ribonucleotide-protein complexes (RNPs). The sgRNA sequences targeting CD33 exon 2 were 5’-TCCATAGCCAGGGCCCCTGT and 5’-GCATGTGACAGGTGAGGCAC.^25^ U937 cells were washed three times in PBS (Gibco 10010-023) and resuspended in complete Nucleofector Kit C (Lonza Biosciences VCA-1004) media (10^6^ cells per transfection) with 5 μL electroporation enhancer (IDT 1075916) and RNPs. Cells were electroporated using a Nucleofector IIb device (Lonza Biosciences) under protocol V-001 and immediately added to a 12 well plate with 1.5 mL complete media and cultured for two weeks.

### Cell Sorting and Flow Cytometry

Edited U937 cells were washed in PBS with 5% heat-inactivated fetal bovine serum (Gibco 10082-147), resuspended at 10^6^ cells/mL and then treated with Human TruStain FcX blocker (BioLegend 422302). Cell sorting was carried out in azide-free buffers; for flow cytometry, 0.02% sodium azide was included in all buffers. Cells were stained with HIM3-4-FITC and P67.6-BV711 for one hour on ice, washed twice with HBSS, then stained with Fixable Viability Dye eFluor780 (Invitrogen 65-0865-18). Cells were resuspended in HBSS (Gibco 24020-117) with 5% heat-inactivated fetal bovine serum (Gibco 10082-147) for sorting. Viable cells were gated using scatter and viability exclusion stain, sorted as either HIM3-4^+^ P67.6^+^, HIM3-4^+^ P67.6^-^, or HIM3-4^-^ P67.6^-^ and collected in complete media. At 48 hours post-sort, cells were split using limiting dilution into a 96 well plate at an average density of 0.5 cells/well and expanded until sufficient cell numbers for analysis were achieved, approximately 8 weeks. Clones were screened by flow cytometry again prior to PCR and sequence analysis.

### PCR Screening and Cloning

Genomic DNA from CRISPR-edited U937 clones was isolated with a DNeasy kit (Qiagen 69506) per manufacturer instructions. A portion of *CD33* was amplified with Q5 High-Fidelity DNA Polymerase (New England BioLabs M0439L) using forward primer 5’-CACAGGAAGCCCTGGAAGCT and reverse primer 5’-GAGCAGGTCAGGTTTTTGGA (Invitrogen). *SIGLEC22P* was amplified similarly with forward primer 5’-GCACCTCAGAGTGGAAGGAC and reverse primer 5’-GAAGGGGTGACTGAGGTACA. Thermocycling parameters were as follows: 98°C 1 min; 98°C 15 s, 66°C 15 s, 72°C 45 s, 32 cycles; 72°C 2 min, 25°C hold. PCR products were separated on a 0.8% agarose-TBE gel, purified using a Monarch gel extraction kit (New England BioLabs T1020L), and sequenced by a commercial company (ACGT, Wheeling, IL). The three missense mutations identified were introduced into a previously described pcDNA3.1-CD33-V5/HIS vector using a QuikChange XL kit (Agilent 200517) with forward primer 5’-GCACTTGCAGCCGGATTTTTGGATCCATAGCCAGGGCC and reverse primer 5’-GGCCCTGGCTATGGATCCAAAAATCCGGCTGCAAGTGC (Invitrogen) to generate the pcDNA3.1-KIR-CD33-V5/HIS vector, transformed into TOP10 E. coli (Invitrogen C404003), isolated using a Plasmid Plus Midiprep Kit (Qiagen 12945) and verified by sequencing (ACGT).^8^

### Gene expression by qPCR

Quantitative PCR (qPCR) was used to quantify expression of total CD33 and D2-CD33 as previously described.^14^ Briefly, primers corresponding to sequences within exons 4 and 5 were used to quantify total *CD33* expression (forward, 5’-TGTTCCACAGAACCCAACAA-3’; reverse, 5’-GGCTGTAACACCAGCTCCTC-3’), as well as primers corresponding to sequences at the exon 1–3 junction and exon 3 to quantify the D2-CD33 isoform (forward, 5’-CCCTGCTGTGGGCAGACTTG-3’; reverse, 5’-GCACCGAGGAGTGAGTAGTCC-3’). PCR was conducted using an initial 2 min incubation at 95°, followed by cycles of 10 s at 95°C, 20 s at 60°C, and 20 s at 72°C. The 20 μl reactions contained 1 μM each primer, 1X PerfeCTa SYBR Green Super Mix (Quanta Biosciences), and 20 ng of cDNA. Experimental samples were amplified in parallel with serially diluted standards that were generated by PCR of cDNA using the indicated primers, followed by purification and quantitation by UV absorbance. Results from samples were compared relative to the standard curve to calculate copy number in each sample. Assays were performed in triplicate and normalized to expression of ribosomal protein L32 (*RPL32*) as the housekeeping gene.^14, 45^

### HEK293 Transfection

HEK293 cells were seeded at approximately 70% confluency 24 hours before transfection. Cells were then transfected with Lipofectamine 3000 with Plus Reagent (Invitrogen L3000001) per manufacturer instructions, 250 ng plasmid per well in 8 well glass chamber slides (MatTek CCS-8) for immunofluorescence or 1000 ng per well in 12 well plates (Corning 3513) for flow cytometry. Cells were transfected with either the previously described wild-type CD33 vector (pcDNA3.1-CD33-V5/HIS), pcDNA3.1-KIRCD33-V5/HIS, or no vector control. Cells were incubated for 24 hours before analysis by flow cytometry or immunofluorescence and confocal microscopy.

### Confocal Immunofluorescence Microscopy

Transfected HEK293 cells were fixed with 10% neutral buffered formalin (Fisher Scientific SF100-4) for 30 minutes then blocked and permeabilized for 30 minutes with 10% goat serum (Sigma S26-LITER), 0.1% Triton X-100 (Fisher Scientific BP151-500) in PBS (Fisher BioReagents BP665-1). Primary and secondary antibodies were diluted in the same blocking and permeabilization buffer and incubated at room temperature for 90 minutes. Cells were washed three times in blocking and permeabilization buffer between primary and secondary antibodies, and three times in PBS prior to coverslip mounting with Prolong Glass with NucBlue mounting media (Invitrogen P36981) and high-tolerance No. 1.5 coverglass (ThorLabs CG15KH1). Images were acquired using a Nikon A1R HD inverted confocal microscope with a 60X oil objective and NIS Elements AR software.

### Statistical analyses

Analyses were performed using GraphPad Prism 8.4.2. Gene expression data were analyzed by one-way ANOVA followed by Dunnett’s multiple comparisons to unedited U937 cells.

## Acknowledgements

This work was supported by RF1AG05971701S1 and R21AG068370 (SE). We would like to thank Yuriko Katsumata for bioinformatics support, Ann M. Stowe for flow cytometry support, the University of Kentucky Light Microscopy Core Facility for help with image acquisition and processing, and the University of Kentucky Flow Cytometry and Immune Monitoring Core Facility, supported in part by the Office of the Vice President for Research, the Markey Cancer Center and an NCI Center Core Support Grant (P30 CA177558).

## Authorship Contributions

BCS conducted all experiments. BCS and SE designed experiments, analyzed data, and drafted the manuscript.

